# binomialRF: Interpretable combinatoric efficiency of random forests to identify biomarker interactions

**DOI:** 10.1101/681973

**Authors:** Samir Rachid Zaim, Colleen Kenost, Joanne Berghout, Wesley Chiu, Liam Wilson, Hao Helen Zhang, Yves A. Lussier

## Abstract

**Background:** In this era of data science-driven bioinformatics, machine learning research has focused on feature selection as users want more interpretation and post-hoc analyses for biomarker detection. However, when there are more features (i.e., transcript) than samples (i.e., mice or human samples) in a study, this poses major statistical challenges in biomarker detection tasks as traditional statistical techniques are underpowered in high dimension. Second and third order interactions of these features pose a substantial combinatoric dimensional challenge. In computational biology, random forest^1^ (**RF**) classifiers are widely used^2–7^ due to their flexibility, powerful performance, and robustness to “P predictors ≫ *subjects N*” difficulties and their ability to rank features. We propose binomialRF, a feature selection technique in RFs that provides an alternative interpretation for features using a correlated binomial distribution and scales efficiently to analyze multiway interactions.

**Methods:** binomialRF treats each tree in a RF as a correlated but exchangeable binary trial. It determines importance by constructing a test statistic based on a feature’s selection frequency to compute its rank, nominal p-value, and multiplicity-adjusted q-value using a one-sided hypothesis test with a correlated binomial distribution. A distributional adjustment addresses the co-dependencies among trees as these trees subsample from the same dataset. The proposed algorithm efficiently identifies multiway nonlinear interactions by generalizing the test statistic to count sub-trees.

**Results:** In simulations and in the Madelon benchmark datasets studies, binomialRF showed computational gains (up to 30 to 600 times faster) while maintaining competitive variable precision and recall in identifying biomarkers’ main effects and interactions. In two clinical studies, the binomialRF algorithm prioritizes previously-published relevant pathological molecular mechanisms (features) with high classification precision and recall using features alone, as well as with their statistical interactions alone.

**Conclusion:** binomialRF extends upon previous methods for identifying interpretable features in RFs and brings them together under a correlated binomial distribution to create an efficient hypothesis testing algorithm that identifies biomarkers’ main effects and interactions. Preliminary results in simulations demonstrate computational gains while retaining competitive model selection and classification accuracies. Future work will extend this framework to incorporate ontologies that provide path-way-level feature selection from gene expression input data.

**Availability:** Github: https://github.com/SamirRachidZaim/binomialRF

**Supplementary information:** Supplementary analyses and results are available at https://github.com/SamirRachidZaim/binomialRF_simulationStudy

## 1 Introduction

Recent advances in machine learning and data science tools have led to a revamped effort for improving clinical decision-making anchored in genomic data analysis and biomarker detection. However, despite these novel advances, random forests (**RFs**) [1] remain a widely popular machine learning algorithm choice in genomics given their ability to i) accurately predict phenotypes using genomic data and ii) identify relevant genes and gene products used for predicting the phenotype. Literature over the past twenty years has demonstrated [2–9] their broad success in being able to robustly handle the “*P* ≫ N” issue where there are more predictors or features “***P***” (i.e., genes) than there are human subjects “*N*” while maintaining competitive predictive and gene selection abilities. However, the translational utility of random forests has not been fully understood as they are often viewed as “black box” algorithms by physicians and geneticists. Therefore, a substantial effort over the past decade has focused around “feature selection” in random forests [5, 6, 10–14] to better provide explanatory power of these models and to identify important genes and gene products in classification models. **Table 1** describes methods of existing feature selection commonly used in random forests as either permutation-type measures of importance, heuristic rankings without formal decision boundaries (i.e., no p-values) or a combination of both. The bioinformatics community have been widely using these feature selection approaches in biomarker discovery [5]. Unfortunately, these techniques do not easily scale computationally nor memory-wise for identifying molecular interactions, seriously limiting their translational utility in medicine and increasing the complexity of their implementation in distributed computing. In addition, there is an increasing consensus among clinicians and machine learning experts that ethical and safe translation of machine learned algorithms for high stake clinical decisions should be interpretable and explainable [15–18].

**Table 1.**
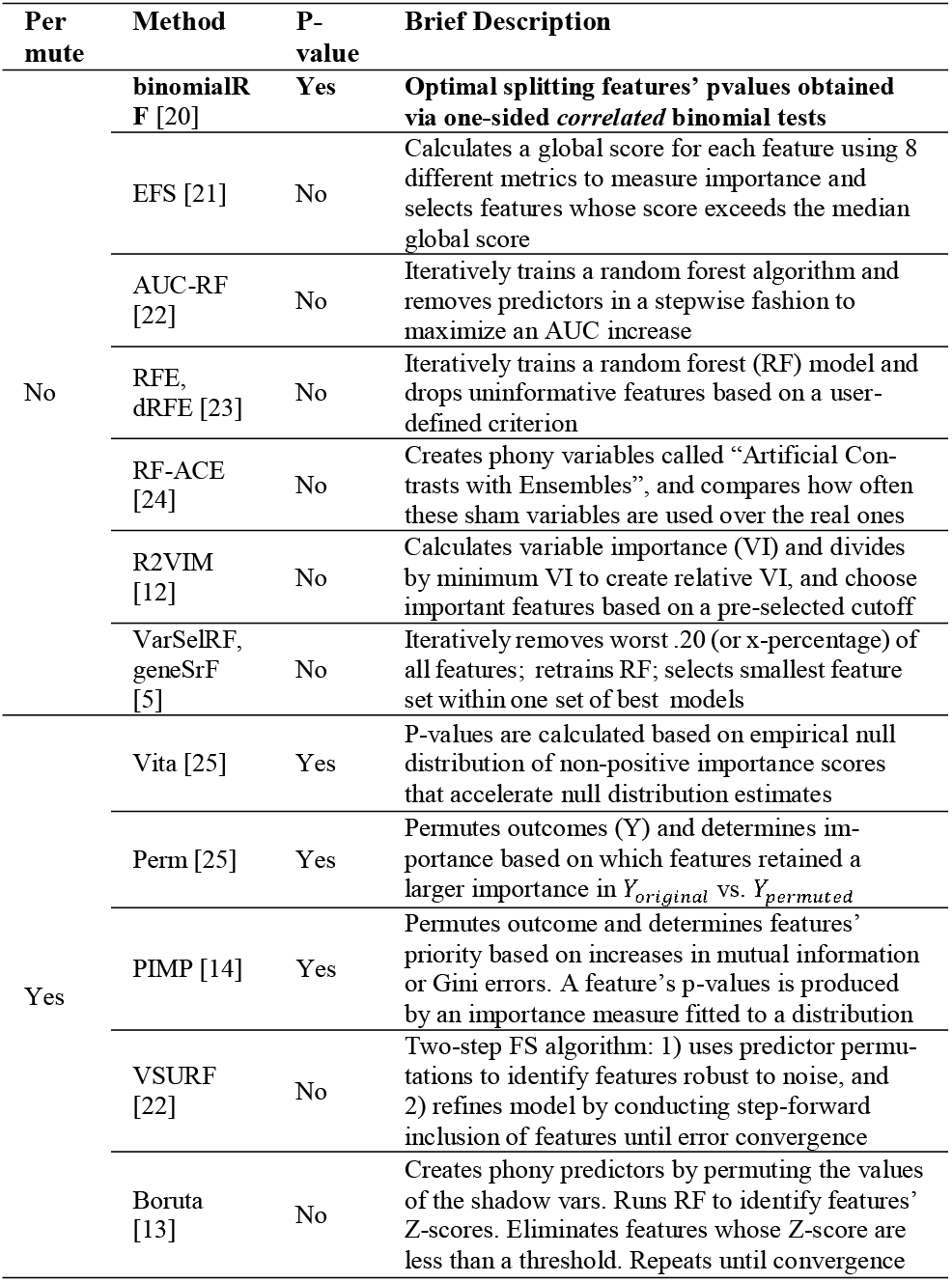
Random forest feature selection methods and their permutation requirements. Absence of permutations generally decreases substantially computing time. P values provide explicit ranking of features, which enables objective feature thresholding.

To address these needs, we propose the ***binomialRF*** feature selection algorithm, a wrapper feature selection algorithm that identifies significant genes and gene sets in a memory-efficient, scalable fashion, with explicit features for biologists and clinicians. Building upon the ‘inclusion frequency”[19] feature ranking, binomialRF formalizes this concept into a binomial probabilistic framework to measure feature importance and extends to identify K-way nonlinear interactions among gene sets. BinomialRF and its extension for model averaging are presented in **Section 2**, while the extension to interaction selection is discussed in **Section 3**. Theoretical computational complexity is presented in **Section 4**, while applications in numerical analyses and case studies evaluate its utility in **Sections 5** and **6**. The discussion, limitations, and concluding sections are presented in **Sections 7** and **8**.

## 2 Methods

We propose a new method for feature selection in random forests, **binomialRF**, which extends and generalizes the “inclusion frequency” strategy to rank features [19] by modeling variable splits at the root of each tree, ***T_Z_***, as a random variable in a stochastic binomial process. This is used to develop a hypothesis-based procedure to model and determine significant features. In the literature, there are a number of existing powerful feature selection algorithms in RF algorithms (see **Table 1**). However, this work proposes an alternative feature selection method using a binomial framework and demonstrates its operating characteristics in comparison to existing technology. **Table 1** illustrates the advantages of the proposed binomialRF as it is both p-value-based and permutation-free, features not identified in our review of literature.

### 2.1 binomialRF notation and information gain from tree splits

Given a dataset, we denote the input information by ***X***, which is comprised of ***N*** subjects (usually < 1,000) and ***P* features (**genes in the genome; usually *P* ≈ 25,000 expressed genes). Genomics data typically represent the “high-dimensional” scenario, where the number of features is much larger than the sample size ***N*** (e.g., “***P*** ≫ *N*”). In the context of binary classification, we denote the outcome variable by ***Y***, which differentiates the case and control groups (i.e., “healthy” vs. “tumor” tissue samples). Random Forests **(RF**) are ensemble learning methods that train a collection of randomized decision trees and construct the decision rule based on combining ***V*** individual trees. We denote a random forest as ***RF*** = {***T*_1_**, …, ***T_V_***}. Each individual decision tree, ***T_z_*** (z = 1, …, ***V***), is trained by using a random subset of the data and features. This randomization encourages a diverse set of trees and allows each individual tree to make predictions across a variety of features and cases. Each tree only sees ***m*** < *P* features in the root when it determines the first optimal feature for splitting the data into two subgroups. The parameter, ***m***, is a user-determined input in the random forest algorithm with default values set usually to either 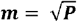 or 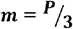. ***F_j,z_*** denotes the random variable measuring whether feature *X_j_* is selected as the splitting variable for tree *T_z_*’s root (**Equation 1**):

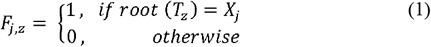

This results in *F_j,z_* following a Bernoulli random variable, *F_j,z_* ~ *Bern*(*p_root_*). In **binomialRF**, to test whether the feature *X_j_* is significant in predicting the outcome **Y**, we build a test statistic 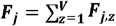 to the the null hypothesis of no feature being significant. One would expect that the probability of selecting a feature *X_j_* would be equal to that of every other feature *X_i_*. Therefore, under the null hypothesis, *p_root_* is constant across all features and trees. Since trees are not independent as they are sampling the same data, *F_j_* follow a ***correlated* binomial distribution** that accounts for the tree-to-tree sampling co-dependencies (**Figure 2**). The following sections will describe combining the probabilistic framework (2.3), the tree-to-tree sampling co-dependency adjustment (2.4), and the test for significance (2.5).

### 2.2 Optimal splitting variable and decision trees

Consider a decision tree, *T_z_*, in a random forest (**Figure 1**). At the top-most “root” node, *m* features are randomly subsampled from the set of *P* features, and the optimal splitting variable, *X_opt_*, is selected as the best feature for separating two classes. Formally, this is stated in **Equation 2.**

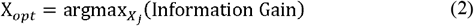

**Fig. 1.**
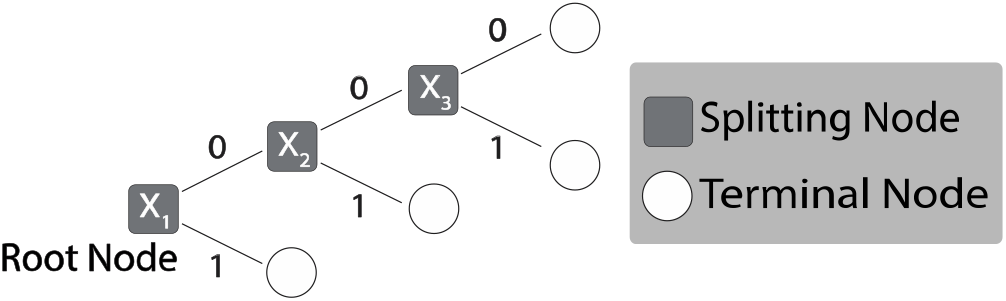
Decision tree and node variables. In the binary split decision tree, *X*_1_ is the optimal splitting feature at the root of the tree, and 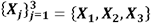 is the optimal splitting sequence that indicates a potential *X*_1_ ⊗ *X*_2_ ⊗ *X*_3_ 3-way interactions, where the symbol “⊗” denotes interactions.

**Fig. 2.**
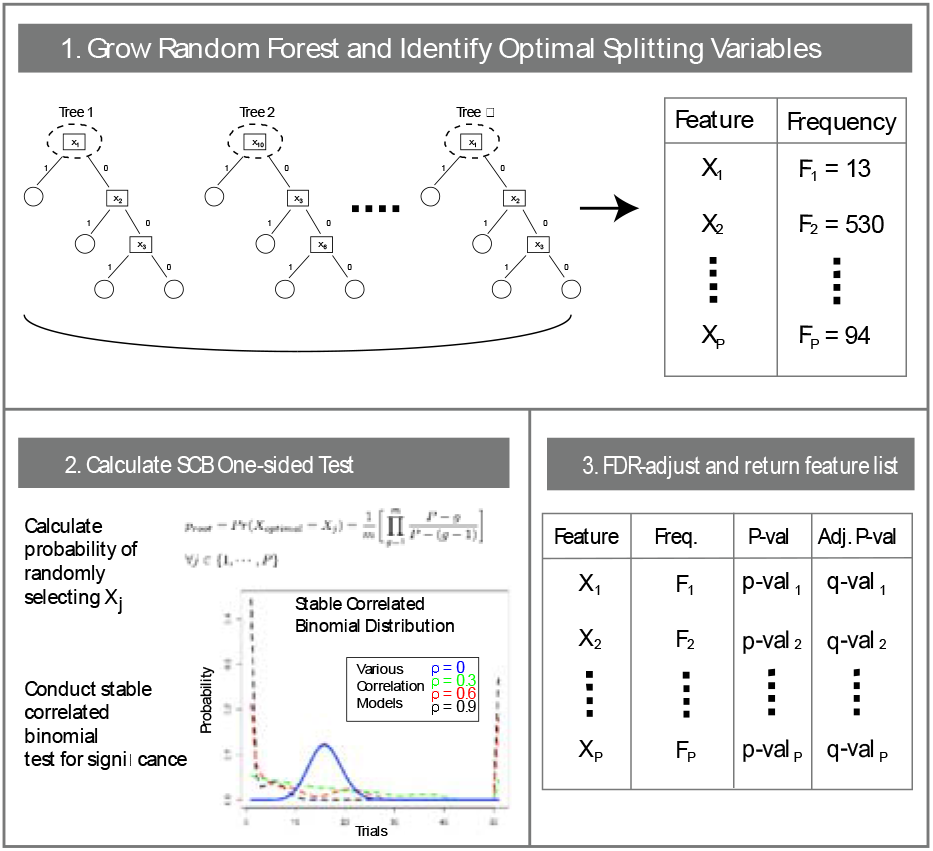
The binomialRF feature selection algorithm. The binomialRF algorithm is a feature selection technique in random forests (**RF**) that treats each tree as a stochastic binomial process and determines whether a feature is selected more often than by random chance as the optimal splitting variable, using a top-bottom sampling without replacement scheme. The main effects algorithm identifies whether the optimal splitting variables at the root of each tree are selected at random or whether certain features are selected with significantly higher frequencies. The interaction-screening extension is detailed in Section 3. *Legend*: *T_z_* = *z^th^* tree in random forest; *X_j_* = feature j; *F_j_* = the observed frequency of selecting *X_j_*; Pr = probability; *P* = number of (#) of features; *V*= # of trees in a RF; m = user parameter to limit *P*; *g* = index of the product.

Focusing on the root, under a null hypothesis, each feature has the same probability of being selected as the optimal root splitting feature, denoted by *p_root_* =Pr(*X_opt_* = *X_j_*) ∀ *j* ∈ {1, …, *P*}. The random variable *F_j,z_* (shown in **Equation 1**) is an indicator variable that tracks if *X_j_* is selected as the optimal variable for the root at tree *T_z_*. *F_j,z_* is a Bernoulli random variable, *F_j,z_* ~ *Bern*(*p_root_*). If all trees were independent, summing across trees yields 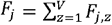 (a binomial random variable). However, trees are not entirely independent since the sampling process creates a co-dependency or correlation across trees.

### 2.3 Adjusting for tree-to-tree co-dependencies

Each tree in a RF samples *n* ⊂ *N* observations either by subsampling or bootstrapping, which creates a tree-to-tree sampling co-dependency, denoted as ***ρ***. In subsampling, the co-dependency between trees is exactly 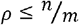, whereas in bootstrapping, the co-dependency is bounded above, i.e., 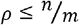. Therefore, in all cases, 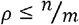 provides a conservative upper bound on the co-dependency between trees. This upper bound adjusts for this tree-to-tree sampling co-dependency. Since the number of sampled cases is determined by the user as a RF parameter, the tree-to-tree co-dependency is known and does not require any estimations. Kuk and Witt both developed a generalization of the family of distributions for exchangeable binary data [26, 27] by adding an extra parameter to model for correlation or association between binary trials when the correlation/association parameter is known. We model this co-dependency among trees by introducing either Kuk’s or Witt’s generalized correlation adjustment in the *correlbinom* R package [26], which is incorporated into the binomialRF model.

### 2.4 Calculating significance of main RF features

At each *T_z_*, *m* < *P* features are subsampled resulting in a probability, *p_root_*, of *X_j_* being selected by a tree, *T_z_*, as shown in **Equation 3**:

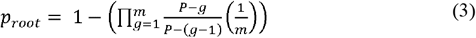

Using **Equation 3**, we can calculate whether *X_j_* provides a statistically significant information gain to discriminate among classes if *F_j_* exceeds the critical value *Q_α,V,p_*, (where *Q_α,V,p_* is the 1 − *α*^th^ quantile of a correlated binomial distribution with *V* trials, *p* is the probability of success, and *ρ* correlation). For multiple hypothesis tests, we adjust our procedure for multiplicity using Benjamini-Yekutieli (BY)[28] false discovery rate.

### 2.5 Calculating significance of RF feature interactions

In classical linear models when detecting 2-way interactions, interactions are included in a multiplicative fashion and treated as separate features with their own linear coefficients. Here, we denote ***X_i_*** ⊗ ***X_j_* as an interaction** between features *X_i_* and *X_j_*. One condition imposed in mathematical interaction selection is strong heredity which states that if the interaction *X_i_* ⊗ *X_j_* is included in the model, then their main effects *X_i_* and *X_j_* must be included. Similarly, under weak heredity, at least one of the two main effects must be included in the model if their interaction term is included. In the context of linear models, several existing methods have been proposed to select interactions and studied in terms of their feasibility and utility [29, 30]. Tree-based methods uniquely bypass these conditions as strong heredity hierarchy is automatically induced resulting from the binary split tree’s structure. As Friedman explains, trees naturally identify interactions based on their sequential, conditional splitting process [31]. This “greedy” search strategy reduces the space from all possible, 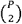 interactions, to only those selected by trees, greatly reducing computational cost and inefficiencies in identifying interactions. We extend previous work in using trees to identify interactions [19, 31] by generalizing the binomialRF to model interactions by considering pairs or sets of sequential splits as random variables and modeling them with the appropriate test statistic and hypothesis test.

To modify the binomialRF algorithm to search for 2-way interactions, we add another product term to **Equation 3** denoting the second feature in the interaction set to calculate *p_2-way_* **(Eq. 4).**

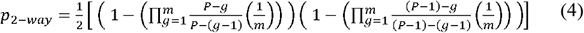

Since we are interested in selecting interactions across variables, if *X_j_* is selected at the root node, then it is no longer available for subsequent selection. Thus, we replace *P* with (*P* − 1). Further, since the interaction can happen two different ways (via the left or right child node), we include a normalizing constant of ½ to account for both ways in which the interaction could occur. **Fig. 3A** illustrates the binomialRF extension to identify 2-way interactions by looking at feature pairs at the root node.

**Fig. 3.**
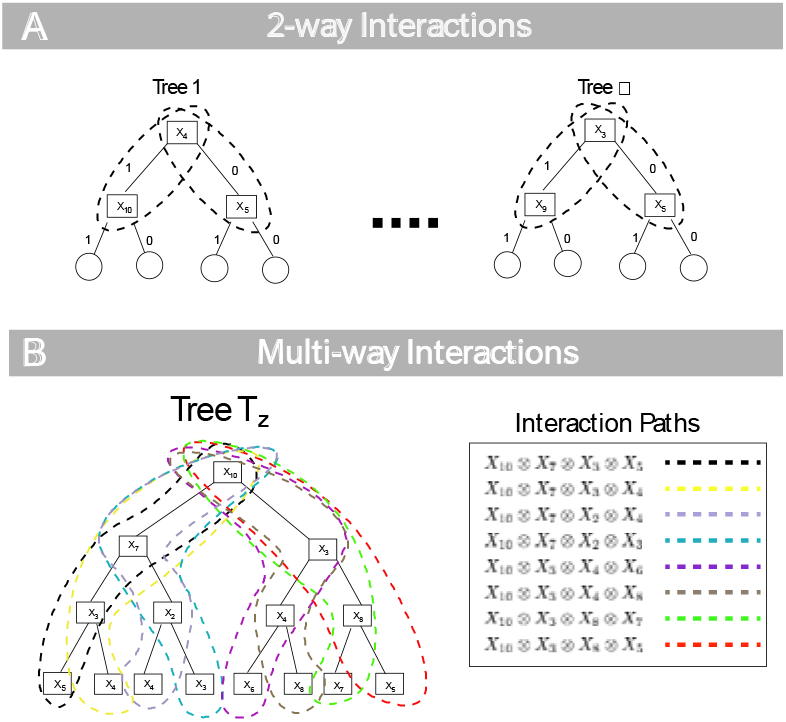
Calculating RF features’ interactions. (A) 2-way Interactions. To extend the binomialRF algorithm for 2-way interaction selection, we define the test statistic which reflects the frequency, *F_ij_* of the pair *X_i_* ⊗ *X_j_* occurring in the random forest. In particular, the probability of an interaction term occurring by random chance is recalculated and normalized by a factor of a half. (B) ***K***-way interactions, ***K*** = 4. Here, we illustrate the tree traversal process to identify all 4-way interactions, ⊗ 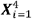, with each color denoting a possible interaction path. The legend on the right shows how each interaction path results in a set of 4-way feature interactions. In general, for any user-desired *K*, the k.binomialRF algorithm traverses the tree via dynamic tree programming to identify all possible paths from the *K*-terminal nodes to the root, where ***K***-terminal nodes are all nodes *K*-steps away from the root node.

To generalize **Equation 4** into multi-way interactions and calculate *p_K−way_*, we first note that for any multi-way interaction of size K in a binary split tree results in at most 2^*K*−l^ terminal nodes. Therefore, there are 2^*K*−l^ possible ways of obtaining the *K*-way interaction (**Fig. 3B**). Thus, the normalizing constant in **Equation 4** is replaced with 2^*K*−l^ in **Equation 5** as a conservative bound on the probability. The product of two terms in **Equation 4** is now expanded to the product of *K* terms (each term representing the probability of selecting one individual feature in the interaction set), and (*P* − 2) is replaced with (*P* − *k*) to account for sampling without replacement, which yields **Equation 5**.

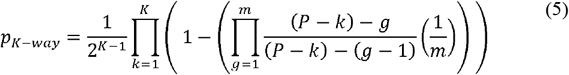

Next, we update the hypothesis test and modify it to identify 2-way interactions for all possible 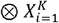 sets.

### 2.6 Evaluation via simulations

To understand the strengths and limitations of the binomialRF feature selection algorithm and to compare its performance with state-of-the-art methods, we conduct a variety of simulations and trials against known benchmark datasets. These simulation scenarios generate logistically-distributed data to mimic binary classification settings in gene expression data using parameters described in **Table 2**: genome size = the dimension of the **X** matrix, a coefficient vector ***β*** that denotes the number of genes seeded linked to the outcome, and the number of trees ***V*** grown in the random forest. The first two parameters are used to generate the design matrix ***X_N×P_***, generate the binary class vector ***Y*** using a logistic regression model, while the final parameter is used to grow the RF.

**Table 2.**
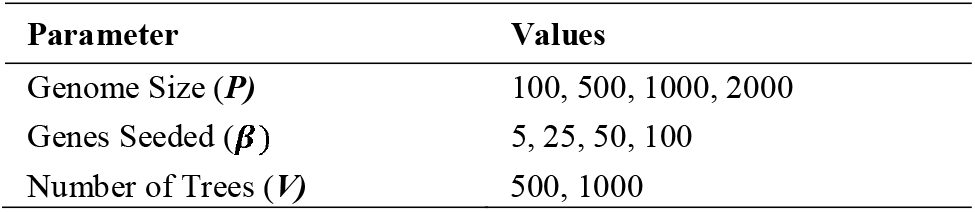
Parameters settings for the simulation study

| Parameter | Values |
| --- | --- |
| Genome Size ($P$) | 100, 500, 1000, 2000 |
| Genes Seeded ($\beta$) | 5, 25, 50, 100 |
| Number of Trees ($V$) | 500, 1000 |

To determine the performance of binomialRF in detecting important interactions, we conduct a simulation study with 30 total features in which we seeded 4 main effects and all 6 possible pairwise interactions. Since the interactions have to be explicitly multiplied in the design matrix, all techniques except binomialRF had a design matrix with all 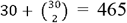 features, and the task was to detect all 6 interactions. Since binomialRF can detect interactions from the original design matrix, we used the original matrix with 30 variables first to identify the main effects and then a second time to identify interactions from main effects.

To evaluate the computational efficiency of binomialRF, we compare the memory requirements and computational time of each method described in **Table 1**. To evaluate classification accuracy, a 0-1 classification loss function is used. Precision (**Eq. 6**), recall (**Eq. 7**), and the Test error (classification error; standard 0/1 loss function) (**Eq. 8**).

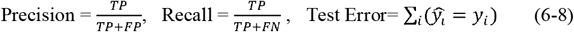

### 2.7 Evaluation in UCI benchmark and TCGA clinical sets

To determine the utility of the binomialRF feature selection algorithm in translational bioinformatics, we conduct a validation study using data from the University of California – Irvine machine learning repository (UCI, hereinafter) and from The Cancer Genome Atlas (TCGA; **Table 3**). The UCI machine learning repository contains over 480 datasets available as benchmarks for machine learning developers to test their algorithms. We present results for all techniques in the Madelon dataset and illustrate their performances using calcification accuracy metrics (cases) presented above in **Equations (6-8)**. Since true variables are not known in these datasets, variable selection accuracies are not calculated.

**Table 3.**
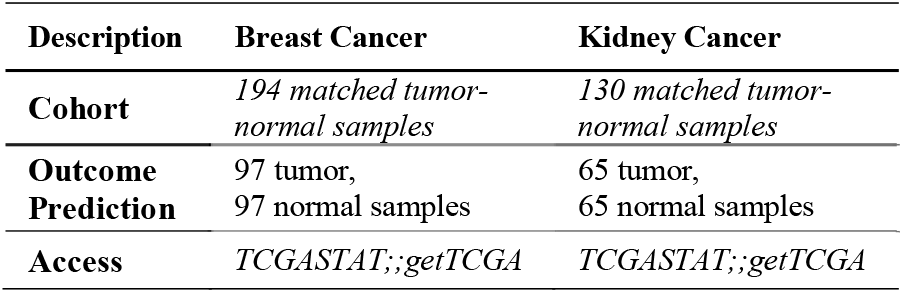
TCGA validation study datasets

We selected the TCGA breast and kidney cancers as two representative datasets with at least 100 matched normal-tumor samples (**Table 3**). The data were downloaded via the R package *TCGA2STAT [32]*, accessed 2020/01, using R.3.5.0. Both RNA sequencing datasets were normalized using RPKM[33]and matched into tumor-normal samples. With many prior studies using the TCGA datasets, our goal was to conduct a binomialRF case study to i) confirm the clinical findings, ii) attain similar prediction performance, and iii) evaluate qualitatively the main effect features and their prioritized interactions. To validate the binomialRF interaction algorithm, we extend the validation of the TCGA datasets <u>*by proposing statistical gene-gene interaction discoveries* and build a classifier from these interactions</u>. We then evaluate their cancer relevance in two ways: (i) trained curators reviewed the literature to identify the involvement of these transcripts in cancer pathophysiology, and (i) a comparison of transcript with the cancer-driving genes of the COSMIC knowledge-base [34].

### 2.8 binomialRF implemented as open source package

The binomialRF R package, wrapping around *randomForest* R package [35], is freely available on GitHub (with simulations, analyses, and results), and has been submitted to CRAN with accompanying documentation and help files (https://github.com/SamirRachidZaim/binomialRF; (https://github.com/SamirRachidZaim/binomialRF_simulationStudy).

## 3 Results

### 3.1 Simulation: computation memory, time, and accuracies

#### Memory storage gains

We use a simple 10-feature simulation. As seen in **Table 4**, it can require as much as 170,000 times less memory to calculate 3-way interactions with binomialRF as compared to classical RF in a moderately large dataset with 1000 variables, which could impact grid computers memory requirements. Note that in linear models, efficient solution paths for 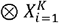 only exist for *K* ∈ {1,2} (LASSO[36] for *K*=1 and RAMP[37] for *K*=2). For *K*>2, to our knowledge, no algorithm guarantees computational efficiency. In RF-based feature selection techniques, the majority of the techniques requires one to explicitly multiply interactions in order to detect them.

**Table 4.**
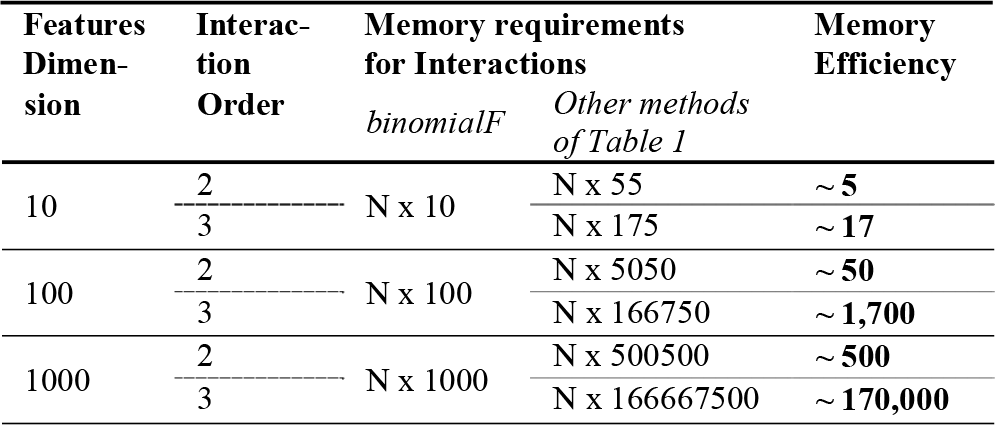
BinomialRF improves the memory requirements by orders of magnitude in 2-way and 3-way interactions when compared to other methods of Table 1. One advantage of the binomialRF algorithm is that it can screen for sets of gene interactions in a memory efficient manner by only requiring a constant-sized matrix whereas the current state of the art requires the predictor matrix to increase in size in a combinatoric fashion to screen for interactions. Memory efficiency is defined by 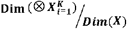 and interaction memory requirements are defined by the number of columns required to map all k-way interactions.

#### Computational time gains

To compare each algorithm’s runtime, we strictly measure the time for the algorithm to produce its feature ranking and omit other portions using the base *system.time* R function. To measure scalability in the predictor space, 500 random forest objects are grown with 500 trees, using simulated genomes sizes 10, 100, and 1000 (**Fig. 4**). **Table 5** illustrates the results for the main effects simulation studies. **Table 5** summarizes all 32 simulation scenarios. **Table 6** shows that in this simulation design, the majority of the techniques were successfully able to identify almost all the interactions (i.e., Recall > .9), and a few were able to do this while also maintaining high precision. Boruta, EFS, binomialRF, and VSURF all achieved at least 70% precision and recall.

**Fig. 4.**
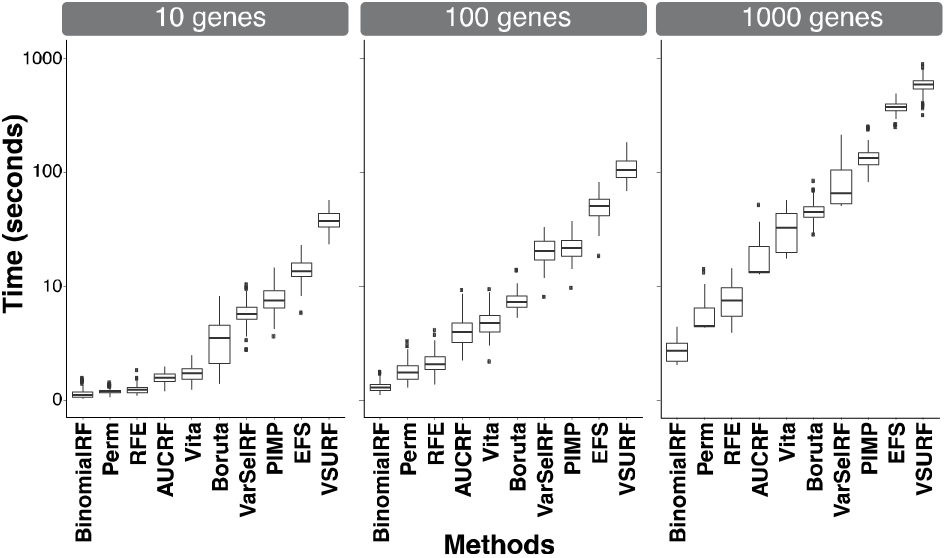
BinomialRF showing substantially improved computational time. The simulation scenarios are detailed in **Section 3.1**, where the length of the coefficient vector, β, varies, but the first five elements are nonzero and P-5 are zero. Rather than measuring accuracy and model identifiability criteria for all the techniques, we only measure the computational runtime of each technique in R. The runtimes are reported in log base 10 (adding +1 to all values to avoid ‘negative’ runtimes), and all simulations were conducted on a 2017 MacBook Pro with 3.1 GHz Intel Core i5 and 16 GB of RAM. All simulations resulted in binomialRF are the fastest. Time presented as log_10_.

**Table 5.**
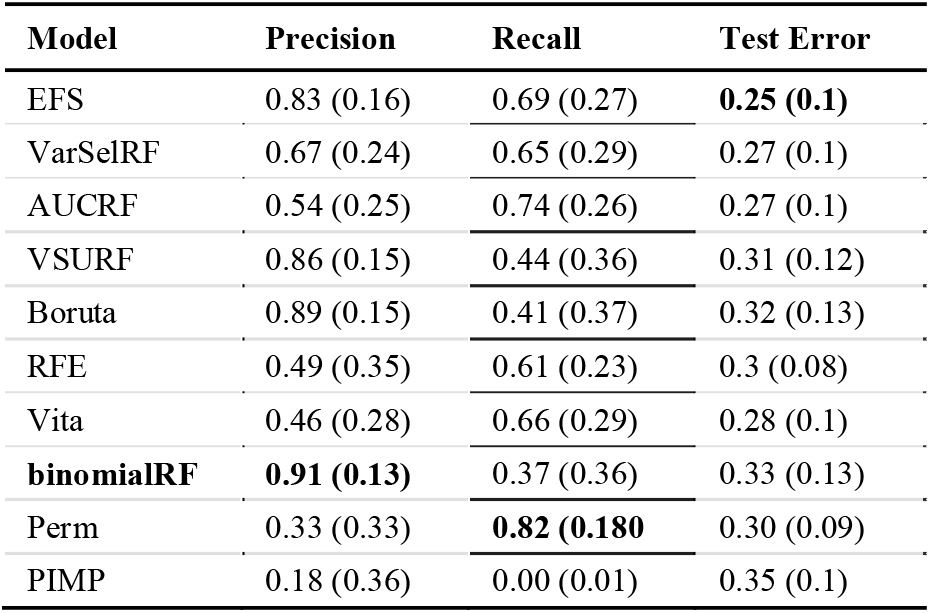
Simulation results of biomarkers. The binomialRF and the algorithms in **Table 1** were tested across a range of simulation scenarios (**Table 2**). Mean (standard deviation) results are shown and ranked according to decreasing harmonic mean of precision and recall of variables. Top accuracies are bolded.

**Table 6.**
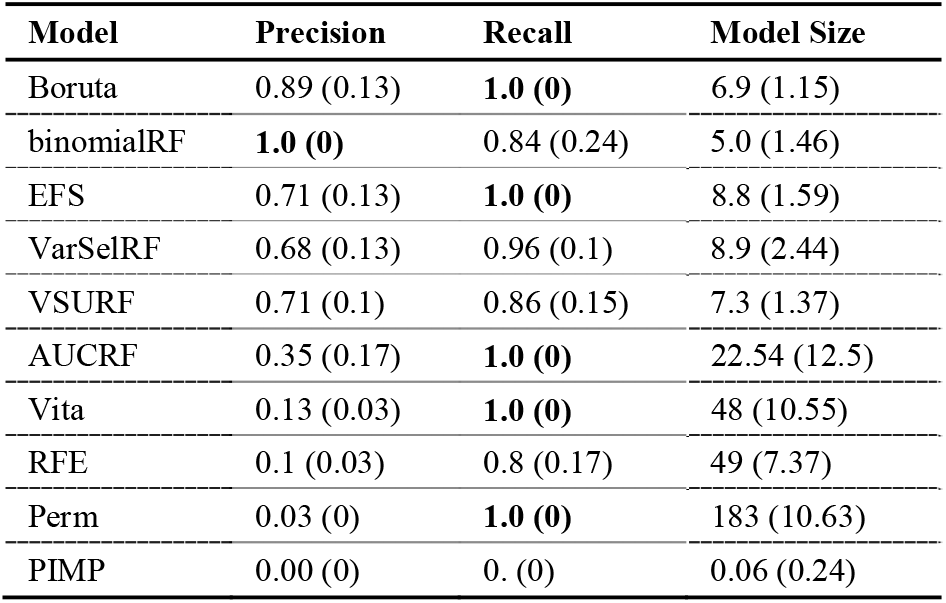
Simulation Results of Biomarker Interactions. Methods are ranked according to decreasing harmonic mean of precision and recall. Top accuracies are bolded.

| Model | Precision | Recall | Model Size |
| --- | --- | --- | --- |
| Boruta | 0.89 (0.13) | **1.0 (0)** | 6.9 (1.15) |
| binomialRF | **1.0 (0)** | 0.84 (0.24) | 5.0 (1.46) |
| EFS | 0.71 (0.13) | **1.0 (0)** | 8.8 (1.59) |
| VarSelRF | 0.68 (0.13) | 0.96 (0.1) | 8.9 (2.44) |
| VSURF | 0.71 (0.1) | 0.86 (0.15) | 7.3 (1.37) |
| AUCRF | 0.35 (0.17) | **1.0 (0)** | 22.54 (12.5) |
| Vita | 0.13 (0.03) | **1.0 (0)** | 48 (10.55) |
| RFE | 0.1 (0.03) | 0.8 (0.17) | 49 (7.37) |
| Perm | 0.03 (0) | **1.0 (0)** | 183 (10.63) |
| PIMP | 0.00 (0) | 0. (0) | 0.06 (0.24) |

### 3.2 UCI ML Benchmark Data Repository

The results for the Madelon dataset show the performance attained by all techniques in a benchmark dataset used to evaluate machine learning algorithms. The results in **Table 7** indicate that all techniques attain a comparable precision and recall, with varying model sizes and run times.

**Table 7.** UCI ML Madelon Dataset Validation. The algorithms in **Table 1** were tested and compared using the Madelon benchmark dataset from UCI (described in Methods). Mean (standard deviation) results are shown and ranked according to decreasing harmonic mean of precision and recall of variables. Top accuracies are bolded.

| Model | Model Size* | Run Time | Precision | Recall |
| --- | --- | --- | --- | --- |
| VarSelRF | 23 (13) | 129 (21) | **0.56 (0.01)** | **0.56 (0.02)** |
| VSURF | 3.5 (1.4) | 321 (267) | **0.56 (0.02)** | **0.56 (0.03)** |
| binomialRF | 17.1 (3.9) | **5.6 (2.2)** | 0.55 (0.02) | 0.55 (0.01) |
| Vita | 13 (5.68) | 1007 (1220) | 0.55 (0.02) | 0.55 (0.02) |
| Boruta | 2 (2) | 139 (45) | 0.54 (0.03) | **0.56 (0.04)** |
| Perm | 240(13) | 269. (329) | 0.56 (0.08) | 0.54 (0.01) |
| AUCRF | 31 (30) | 33 (7.5) | 0.55 (0.04) | 0.54 (0.02) |
| RFE | 81 (4.2) | 20 (1.4) | 0.54 (0.06) | 0.54 (0.01) |
| EFS | 20 (8.3) | 2617 (2126) | 0.53 (0.02) | 0.54 (0.02) |
| PIMP | 1.7 (1.3) | 482 (128) | 0.50 (0.04) | 0.50 (0.01) |

### 3.3 TCGA clinical validations in breast and kidney cancers

**Figure 5** indicates that although the black-box classifier with all >19,000 genes results in a highly accurate classifier (i.e., precision and recall >0.98), that the binomialRF classifiers with only 51 genes in breast cancer and 16 in kidney cancer, respectively, obtained comparable performances. Furthermore, after identifying key statistical interactions (39 in breast, 11 in kidney), we validated their signal by building a classifier exclusively from them with comparable accuracy.

**Fig.5.**
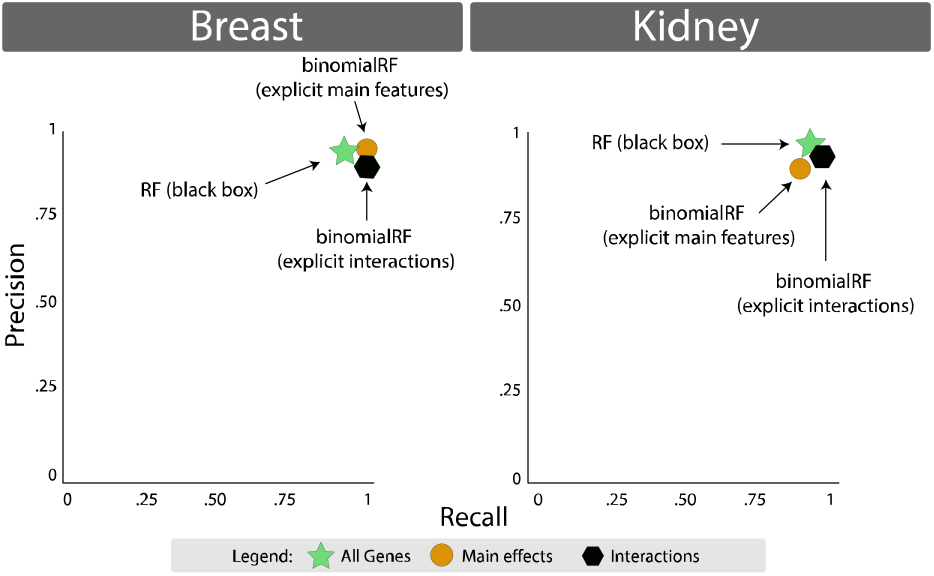
Biomarker accuracies of the TCGA validation study. The TCGA validation study was conducted using breast and kidney cancer datasets, accessed via the R package *TCGA2STAT*. The matched-sample datasets were utilized to determine whether binomialRF could produce an accurate classifier via main effects and interactions. Left, the two binomialRF classifiers (51 identified gene main effects; 39 identified gene-gene interactions) and obtained a classifier as accurate as the original black-box RF model with all ~20,000 genes. Right, the two binomialRF classifiers (16 identified gene main effects; 11 identified gene-gene interactions) obtained a classifier as accurate as the original black-box RF model with all ~20,000 genes.

To validate the identified interactions across both TCGA studies, we constructed networks of their pairwise statistical interactions and assessed whether the log-ratio of the gene expression were distributed differently across tumor and normal samples. **Fig. 6** provides the statistical interaction networks, as well as exemplar cases of a gene-gene interactions in each of the studies. For breast cancer, we present an interaction between SPRY2 and C0L10A1 and for kidney one between TFAP2A and SGPP1. In each study, the two individual genes in isolation are expressed differently across normal-tumor samples indicative of their discrimination power. Further, the log-ratios of both genes show an additional level of statistical signal that is captured from the interaction, suggesting the possibility of biological interaction.

**Fig. 6.**
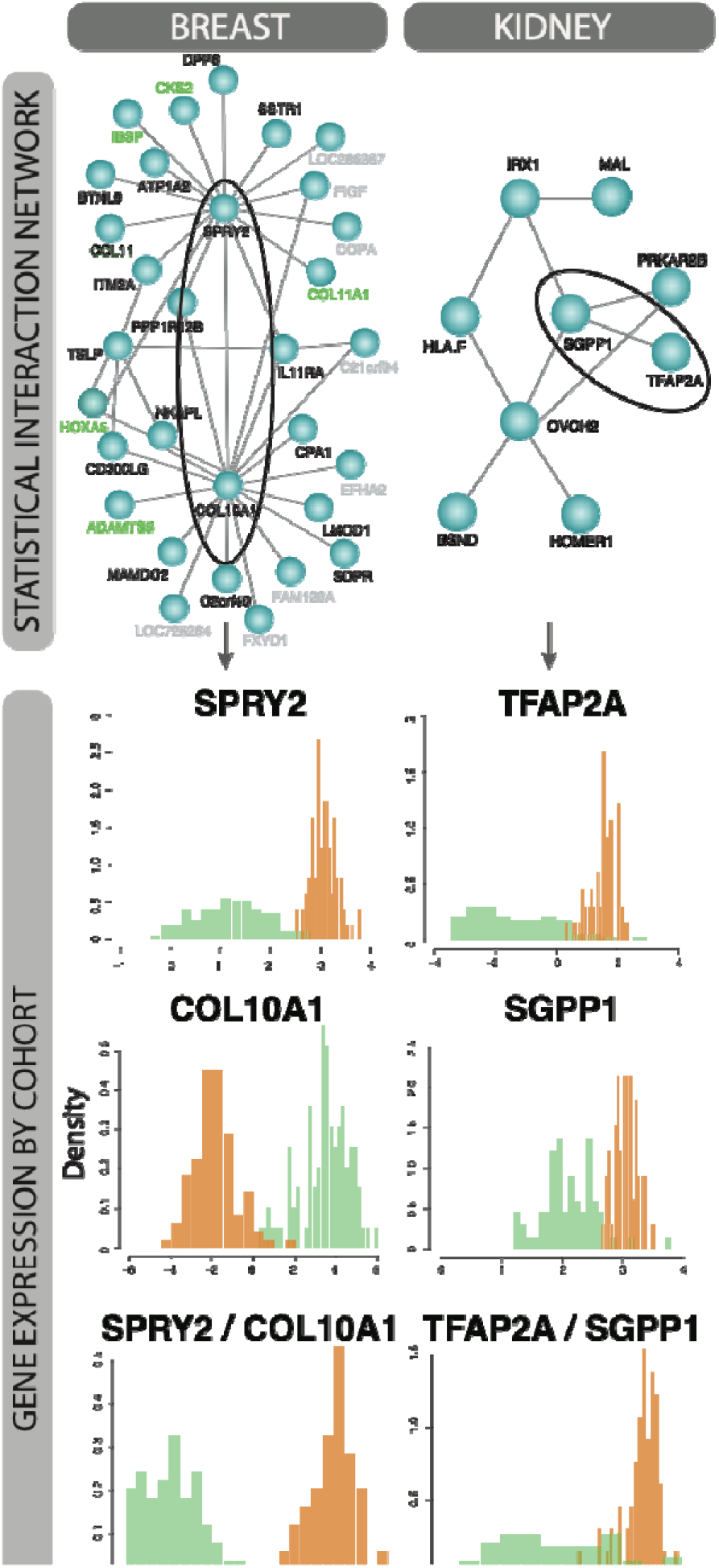
Statistical interactions prioritized by binomialRF in TCGA cancers recapitulate known cancer driver genes. The statistical interaction gene networks (Top) indicate the pairwise biomarker interactions identified by the binomialRF algorithm for the breast (Left) and kidney (Right) cancer datasets. Key features are involved in multiple interactors (super-interactors; e.g., SPRY2; COL10A1). Features names (gene products) found in the literature as associated to cancer pathophysiology are shown in black; those also documented as driving cancer genes in COSMIC are shown in green (Methods); the remainder are grey. Two exemplar statistical interactions (one per dataset) are circled and the log expression of their gene products and of their ratios are shown in the bottom panels. The distribution separation across tumor (green) and normal (orange) cases indicates a potential interaction between these two genes across the cohorts.

## 4 Discussion

### 4.1 Numerical studies, RF-based feature selection techniques, efficiency gains, and interactions

The averaged results across all 32 simulation designs are presented in **Table 5**, with each category’s best values highlighted in bold. Techniques such as AUCRF and EFS result in the smallest prediction error, showcasing their strength in the prediction task. The permutation resampling strategy attains the highest recall, which provides users a tool to identify gene products that are potentially relevant for a disease. For main effects and for statistical interactions, Boruta, VSURF, and binomialRF algorithms attain the highest precisions (positive predictive value) with reasonable recall. In addition, EFS also performs at reasonable recall and precision for the interactions. BinomialRF distinguished itself with the most optimal memory utilization and runtimes.

Strobl and Zeileis [38] demonstrate that i) the *Gini importance* (measure of entropy) is biased towards predictors with many categories, and ii) that growing more trees inflates anticonservative power estimates. To address (i), we recommend the user evaluates sets of genes according to their baseline expression levels [39]. For the latter (ii), the binomialRF uses *ntree* parameter (number of trees; **Table 2**) to calculate a conservative cumulative distribution function (cdf) rather than calculating an anticonservative (**Eq. 1**), which mitigates the possibility of overtraining. Our simulations were ran using 500 and 1000 trees with no visible differences across results (**Tables 2 and 5**). We ran five additional simulations (seeding 5/100 genes) using 100, 200, 500, 1000, and 2000 trees to determine the effect of growing more trees. The median results indicate that as the number of trees increases, the metrics tend to converge (data not shown), indicating a stability in the number of trees. To test false positive results, we ran three additional simulations with absence of informative features in the simulation (genes seeded =0; data not shown), indicating that in absence of informative variables, the binomialRF produces a type I error ranging between 0.5 - 2%. Future simulations will explore artificial datasets with main effects in absence of interactions to quantify type I errors.

There are other complementary efforts to improve the efficiency of random forests. Studies [40–43] focus on subspace sampling methods, reducing the search, and ensuring diversity among the features or cases sampled to make the node-splitting process more efficient, rather than biomarker discoveries. Other sets of techniques (such as [44]) gain efficiency by modifying the learning process. These methods are independent of feature selection and could be combined with any method from **Table 1** to further improve RF efficiencies.

binomialRF proposes an automated combinatoric memory reduction in the original predictor matrix (Table 4), while other methods from Table 1 generally require rate-limiting and memory consuming userdefined explicit interactions by multiplying the interactions. Alternatively, using recursive partitioning to identify interactions dates back to 2004 in random forests [45], and as recent as [19] with Bayesian regression trees and joint-feature inclusion frequencies. Many studies identify sets of conditional or sequential splits, while other strategies (i.e., [46]) measure their effect in prediction error. binomialRF automatically models these sequential split frequencies into a hypothesis testing framework using a generalization of the binomial distribution that adjusts for tree-totree data co-dependency. This contribution provides an alternative p-value-based strategy to explicitly rank feature interactions in any order with the binomialRF, using a simple modification of a user-determined parameter, *k*. Future work will aim to refine and polish interaction detection within the binomialRF framework and extend the preliminary results and techniques.

### 4.2 Moving towards interpretable, white-box algorithms

In recent years, there have been substantial efforts to develop more human-interpretable machine learning tools in response to the ethical and safety concerns of using ‘blackbox’ algorithms in medicine [15] or in high stake decisions[16]. A perspective on *Nature Machine Intelligence* [16], the Explainable Machine Learning Challenge in 2018 [47], and other initiatives serve as reminders of the ethical advantages of using interpretable white-box models over blackbox. Novel software packages and methods (i.e., [48, 49]) bring elements of ensemble learning and RFs into the linear model space to combine the high accuracy of ensemble learners with interpretability of generalized linear models. Other initiatives such as the *iml* R package [49] provide post-hoc interpretability tools for blackbox algorithms or provide model-agnostic strategies “to *trust* and act on predictions” [50]. These white-box efforts are converging towards producing more explanatory power that improve ethical and safe decision making. Feature selection methods also improve the transparency of machine learning methods. Further, there is a need to develop algorithms that can better illustrate how they have identified features, and how these algorithms rank features. Among feature selection techniques, binomialRF provides more explicit features and their interactions than conventional RF as well as a prioritization statistic. This differs from the majority of other feature selection methods that have been developed for RF, as they do not provide a prioritization among features (Table 1; p-value= no). For the remainder that provide p-values, they require memory intensive and time-consuming permutation tests. While BinomialRF is permutation free, it nonetheless is not fully a white box in that the precise decision tree is not explicit. Future work will extend binomialRF to develop stronger interpretation and visualizing tools (i.e., **Figure 6**) of these trees (e.g., median accuracy tree among competing ones with the best prioritized features and interactions).

As recent work by our lab and others have shown, there is a subspace of genomic classifiers and biomarker detection anchored in pathways and ontologies[51–53] that has yielded promising results in biomarker detection using *a priori* defined gene sets (i.e., GO[54]). Hsueh et al. have explored the subdomain of ontology-anchored gene expression classifiers in random forests [55] They also discuss alternate statistical techniques available for geneset analyses and paved the way towards RF-based geneset analysis. In future work, we will direct our efforts along this path and extend binomialRF to incorporate gene set-anchored feature selection algorithms that explore pathway interactions.

## 5. Conclusion

We propose a new feature selection method for exploring feature interactions in random forests, binomialRF, and shown in simulations its sparse memory usage and substantially improved computational run time as compared to previous methods. It was also among the top four most accurate (precision, recall) among ten techniques across large scale simulations and benchmark datasets. In addition, in clinical datasets, the prioritized interaction classifiers attain high performance with less than 1% of the features and produce pathophysiologically relevant features (evaluated via curation and external reference standards). We have released an open source package in R on GitHub and have submitted it to the CRAN (R archive) for consideration.

Machine learning algorithms are increasingly required to explain their predictions and features in human-interpretable form for high stake decision making; therefore, more methods need to be developed that provide explicit white-box-style classifiers with the high accuracy rates otherwise observed in conventional blackbox-style algorithms (e.g., random forests). Among feature selection methods designed for random forests, binomialRF proves to be more efficient and as accurate for exploring high order interactions between biomolecular features as compared to ten published methods. This increased efficiency for exploring complexity may contribute to improving therapeutic decision making, which may address existing machine learning gaps in precision medicine.

## List of acronyms and symbols

⊗: Symbol denoting interaction
BY: Benjamini Yekutieli adjustment
CDF: cumulative distribution function
GO: Gene Ontology
RF: random forest
UCI: University of California – Irvine
TCGA: The Cancer Genome Atlas

## Acknowledgements

The authors would like to acknowledge the University of Arizona’s High-Performance Computing (HPC) for providing the space and computing hours to conduct our simulation studies and analyses.

## Author Contributions

SRZ conducted all the analyses in R; HHZ contributed to the statistical framework and analysis; all authors contributed to the evaluation and interpretation of the study; SRZ, JB, WC, LW, and CK contributed to the figures; SRZ, JB, WC, LW, and CK contributed to the tables; JB, WC, LW contributed to the evaluation of the clinical studies; SRZ, HHZ, CK, and YAL contributed to the writing of the manuscript; all authors read and approved the final manuscript.

## References

1. Breiman, L., Random forests. Machine learning, 2001. 45(1): p. 5–32.

2. Chen, X. and H. Ishwaran, Random forests for genomic data analysis. Genomics, 2012. 99(6): p. 323–329.

3. Bienkowska, J.R., et al., Convergent Random Forest predictor: methodology for predicting drug response from genome-scale data applied to anti-TNF response. Genomics, 2009. 94(6): p. 423–432.

4. Boulesteix, A.L., et al., Overview of random forest methodology and practical guidance with emphasis on computational biology and bioinformatics. Wiley Interdisciplinary Reviews: Data Mining and Knowledge Discovery, 2012. 2(6): p. 493–507.

5. Diaz-Uriarte, R., GeneSrF and varSelRF: a web-based tool and R package for gene selection and classification using random forest. BMC bioinformatics, 2007. 8(1): p. 328.

6. Díaz-Uriarte, R. and S.A. De Andres, Gene selection and classification of microarray data using random forest. BMC bioinformatics, 2006. 7(1): p. 3.

7. Goldstein, B.A., et al., An application of Random Forests to a genome-wide association dataset: methodological considerations & new findings. BMC genetics, 2010. 11(1): p. 49.

8. Izmirlian, G., Application of the random forest classification algorithm to a SELDI□TOF proteomics study in the setting of a cancer prevention trial. Annals of the New York Academy of Sciences, 2004. 1020(1): p. 154–174.

9. Jiang, P., et al., MiPred: classification of real and pseudo microRNA precursors using random forest prediction model with combined features. Nucleic acids research, 2007. 35(suppl_2): p. W339–W344.

10. Archer, K.J. and R.V. Kimes, Empirical characterization of random forest variable importance measures. Computational Statistics & Data Analysis, 2008. 52(4): p. 2249–2260.

11. Genuer, R., J.-M. Poggi, and C. Tuleau-Malot, VSURF: an R package for variable selection using random forests. The R Journal, 2015. 7(2): p. 19–33.

12. Szymczak, S., et al., r2VIM: A new variable selection method for random forests in genome-wide association studies. BioData mining, 2016. 9(1): p. 7.

13. Kursa, M.B. and W.R. Rudnicki, Feature selection with the Boruta package. J Stat Softw, 2010. 36(11): p. 1–13.

14. Altmann, A., et al., Permutation importance: a corrected feature importance measure. Bioinformatics, 2010. 26(10): p. 1340–1347.

15. Char, D.S., N.H. Shah, and D. Magnus, Implementing machine learning in health care—addressing ethical challenges. The New England journal of medicine, 2018. 378(11): p. 981.

16. Rudin, C., Stop explaining black box machine learning models for high stakes decisions and use interpretable models instead. Nature Machine Intelligence, 2019. 1(5): p. 206–215.

17. Možina, M., J. Žabkar, and I. Bratko, Argument based machine learning. Artificial Intelligence, 2007. 171(10-15): p. 922–937.

18. Watson, D.S., et al., Clinical applications of machine learning algorithms: beyond the black box. Bmj, 2019. 364: p. l886.

19. Chipman, H.A., E.I. George, and R.E. McCulloch, BART: Bayesian additive regression trees. The Annals of Applied Statistics, 2010. 4(1): p. 266–298.

20. Zaim, S.R., et al., binomialRF: Scalable Feature Selection and Screening for Random Forests to Identify Biomarkers and Their Interactions. bioRxiv, 2019: p. 681973.

21. Neumann, U., N. Genze, and D. Heider, EFS: an ensemble feature selection tool implemented as R-package and web-application. BioData mining, 2017. 10(1): p. 21.

22. Calle, M.L., et al., AUC-RF: a new strategy for genomic profiling with random forest. Human heredity, 2011. 72(2): p. 121–132.

23. Nguyen, H.-N. and S.-Y. Ohn. Drfe: Dynamic recursive feature elimination for gene identification based on random forest. in International Conference on Neural Information Processing. 2006. Springer.

24. Tuv, E., et al., Feature selection with ensembles, artificial variables, and redundancy elimination. Journal of Machine Learning Research, 2009. 10(Jul): p. 1341–1366.

25. Degenhardt, F., S. Seifert, and S. Szymczak, Evaluation of variable selection methods for random forests and omics data sets. Briefings in bioinformatics, 2017. 20(2): p. 492–503.

26. Witt, G., A Simple Distribution for the Sum of Correlated, Exchangeable Binary Data. Communications in Statistics-Theory and Methods, 2014. 43(20): p. 4265–4280.

27. Kuk, A.Y., A litter□based approach to risk assessment in developmental toxicity studies via a power family of completely monotone functions. Journal of the Royal Statistical Society: Series C (Applied Statistics), 2004. 53(2): p. 369–386.

28. Benjamini, Y. and D. Yekutieli, The control of the false discovery rate in multiple testing under dependency. The annals of statistics, 2001. 29(4): p. 1165–1188.

29. Nelder, J.A., The selection of terms in response-surface models—how strong is the weak-heredity principle? The American Statistician, 1998. 52(4): p. 315–318.

30. Choi, N.H., W. Li, and J. Zhu, Variable selection with the strong heredity constraint and its oracle property. Journal of the American Statistical Association, 2010. 105(489): p. 354–364.

31. Friedman, J., T. Hastie, and R. Tibshirani, The elements of statistical learning. Vol. 1. 2001: Springer series in statistics New York.

32. Wan, Y.-W., G.I. Allen, and Z. Liu, TCGA2STAT: simple TCGA data access for integrated statistical analysis in R. Bioinformatics, 2016. 32(6): p. 952–954.

33. Wagner, G.P., K. Kin, and V.J. Lynch, Measurement of mRNA abundance using RNA-seq data: RPKM measure is inconsistent among samples. Theory in biosciences, 2012. 131(4): p. 281–285.

34. Bindal, N., et al., COSMIC: the catalogue of somatic mutations in cancer. Genome biology, 2011. 12(1): p. P3.

35. Liaw, A. and M. Wiener, Classification and regression by randomForest. R news, 2002. 2(3): p. 18–22.

36. Tibshirani, R., Regression shrinkage and selection via the lasso. Journal of the Royal Statistical Society: Series B (Methodological), 1996. 58(1): p. 267–288.

37. Hao, N., Y. Feng, and H.H. Zhang, Model selection for high-dimensional quadratic regression via regularization. Journal of the American Statistical Association, 2018. 113(522): p. 615–625.

38. Strobl, C. and A. Zeileis, Danger: High power!–exploring the statistical properties of a test for random forest variable importance. 2008.

39. Li, Q., et al., Interpretation of Omics dynamics in a single subject using local estimates of dispersion between two transcriptomes. bioRxiv, 2019: p. 405332.

40. Wang, Q., et al., An efficient random forests algorithm for high dimensional data classification. Advances in Data Analysis and Classification, 2018. 12(4): p. 953–972.

41. Wu, Q., et al., ForesTexter: an efficient random forest algorithm for imbalanced text categorization. Knowledge-Based Systems, 2014. 67: p. 105–116.

42. Ye, Y., et al., Stratified sampling for feature subspace selection in random forests for high dimensional data. Pattern Recognition, 2013. 46(3): p. 769–787.

43. Sinha, V.Y.K.P.K. and V.Y. Kulkarni. Efficient learning of random forest classifier using disjoint partitioning approach. in Proceedings of the World Congress on Engineering. 2013.

44. Lakshminarayanan, B., D.M. Roy, and Y.W. Teh. Mondrian forests: Efficient online random forests. in Advances in neural information processing systems. 2014.

45. Lunetta, K.L., et al., Screening large-scale association study data: exploiting interactions using random forests. BMC genetics, 2004. 5(1): p. 32.

46. Li, J., et al., Detecting gene-gene interactions using a permutation-based random forest method. BioData mining, 2016. 9(1): p. 14.

47. Rudin, C. and J. Radin, Why Are We Using Black Box Models in AI When We Don’t Need To? A Lesson From An Explainable AI Competition. Harvard Data Science Review, 2019. 1(2).

48. Song, L., P. Langfelder, and S. Horvath, Random generalized linear model: a highly accurate and interpretable ensemble predictor. BMC bioinformatics, 2013. 14(1): p. 5.

49. Molnar, C., G. Casalicchio, and B. Bischl, iml: An R package for interpretable machine learning. Journal of Open Source Software, 2018. 3(26): p. 786.

50. Ribeiro, M.T., S. Singh, and C. Guestrin, Model-agnostic interpretability of machine learning. arXiv preprint arXiv:1606.05386, 2016.

51. Zaim, S.R., et al., Emergence of pathway-level composite biomarkers from converging gene set signals of heterogeneous transcriptomic responses. 2018.

52. Gardeux, V., et al., ‘N-of-1-pathways’ unveils personal deregulated mechanisms from a single pair of RNA-Seq samples: towards precision medicine. Journal of the American Medical Informatics Association, 2014. 21(6): p. 1015–1025.

53. Gardeux, V., et al., A genome-by-environment interaction classifier for precision medicine: personal transcriptome response to rhinovirus identifies children prone to asthma exacerbations. Journal of the American Medical Informatics Association, 2017. 24(6): p. 1116–1126.

54. Ashburner, M., et al., Gene ontology: tool for the unification of biology. Nature genetics, 2000. 25(1): p. 25.

55. Hsueh, H.-M., D.-W. Zhou, and C.-A. Tsai, Random forests-based differential analysis of gene sets for gene expression data. Gene, 2013. 518(1): p. 179–186.

